# Melanocortin receptor agonist melanotan-II microinjected in the nucleus accumbens decreases appetitive and consumptive responding for food

**DOI:** 10.1101/2022.04.13.488209

**Authors:** Nicole Eliason, Lynne Martin, Malcolm J. Low, Amanda L. Sharpe

## Abstract

*Rationale*. Obesity is a major health problem worldwide. An understanding of the factors that drive feeding behaviors is key to the development of pharmaceuticals to decrease appetite and consumption. Proopiomelanocortin (POMC), the melanocortin peptide precursor, is essential in the regulation of body weight and ingestive behaviors. Deletion of POMC or impairment of melanocortin signaling in the brain results in hyperphagic obesity. Neurons in the hypothalamic arcuate nucleus produce POMC and project to many areas including the nucleus accumbens (NAcc), which is well established in the rewarding and reinforcing effects of both food and drugs of abuse.

**Objective:** These studies sought to determine the role of melanocortins in the NAcc on consumption of and motivation to obtain access to standard rodent chow.

**Methods:** Male, C57BL/6J mice were microinjected bilaterally into the NAcc (100 nl/side) with the melanocortin receptor 3/4 agonist melanotan-II (MT-II; 0.1, 0.3, and 1 nmol), and ingestive behaviors were examined in both home cage and operant food self-administration experiments. In addition, the ability of MT-II in the NAcc to produce aversive properties or affect metabolic rate were tested.

**Results:** MT-II injected into the NAcc significantly decreased consumption in both home cage and operant paradigms, and furthermore decreased appetitive responding to gain access to food. There was no development of conditioned taste avoidance or change in metabolic parameters following anorexic doses of MT-II. *Conclusions*. MT-II in the NAcc decreased both the motivation to eat and the amount of food consumed without inducing an aversive state or affecting metabolic rate, suggesting a role for melanocortin signaling in the ventral striatum that is selective for appetite and satiety without affecting metabolism or producing an aversive state.

## INTRODUCTION

Obesity is an imminent threat to public health due to the co-morbidity with diabetes, cancer, heart disease, and depression. Homeostatic regulation of body weight is achieved through balance between calories consumed and metabolic rate of the organism. Consumption of food is driven by both metabolic demand and by rewarding properties of food. An understanding of the neurocircuitry that drives both metabolic and reward components of feeding is critical for the treatment of ingestive disorders and obesity.

Proopiomelanocortin (POMC) peptides are known to be crucial in body weight homeostasis. Deletion of POMC in whole animal or exclusively in the brain creates a phenotype of morbid obesity due to both hyperphagia and a decreased metabolic rate (Smart et al. 2006; Yaswen et al. 1999). Melanocortin peptides, produced in the brain through cleavage of POMC, are an important component of POMC’s role in energy homeostasis as both mice and humans with a defect of the melanocortin receptors exhibit morbid obesity and impaired metabolic rate (Huszar et al. 1997; Ste Marie et al. 2000; Vaisse et al. 1998; Yeo et al. 1998). Early pharmacological studies established a central role in both metabolic rate and food intake for melanocortins through ventricular injection of melanocortin receptor agonists and antagonists (Azzara et al. 2002; Benoit et al. 2000, 2001, 2003; Fan et al. 1997; Olszewski et al. 2001; Thiele et al. 1998; Wirth et al. 2001; Zheng et al. 2010). While POMC neuronal populations may be important for both feeding and metabolic rate, it is possible that they may be regulated by different pathways. In treating obesity it may be more beneficial to decrease feeding without increasing metabolic rate and possible subsequent adverse cardiovascular effects. Thus, understanding the physiological effect (feeding versus metabolic rate) mediated by melanocortins in different brain regions is crucial to therapeutic developments that maximize efficacy while minimizing adverse effects.

POMC expressing neurons are located in the arcuate nucleus of the hypothalamus and the nucleus of the solitary tract in the brainstem. The arcuate nucleus neurons project to many regions of the brain associated with feeding and reward, including other regions of the hypothalamus, the amygdala, the nucleus of the stria terminalis, and the NAcc (Eskay et al. 1979; Chronwall 1985). Previous studies microinjecting melanocortin agonists into the paraventricular nucleus of the hypothalamus (Beckman et al. 2009; Giraudo et al. 1998; Kim et al. 2000; Olszewski et al. 2001; Skibicka and Grill 2009; Wirth et al. 2001) have established the importance of melanocortins in this brain region in both satiety and energy expenditure.

POMC projections to the nucleus accumbens (NAcc) are of interest in feeding studies because of the established role of the NAcc in reward and reinforcement for both natural (food) and drug substances. Injection of melanocortin agonists in the NAcc of rats decreases intake of regular chow, sucrose, and ethanol (Lerma-Cabrera et al., 2012; Pandit et al., 2015). However, other studies microinjecting melanocortin receptor antagonists into the NACC have produced conflicting findings showing that melanocortins are necessary for the rewarding and reinforcing effects of cocaine (Hsu et al. 2005) but have no role in food consumption (Kask and Schioth 2000, but also see Davis et al. 2011). Differences in the food reinforcer used (sucrose or chow), operant versus free-feeding, and other parameters have varied among these studies—potentially adding to the dissonance. In addition, these measures were largely focused on consumption with no measures of reward independent of intake.

The objective of our study was to investigate the role of melanocortin receptor agonists in the NAcc on appetitive (motivation to gain access to food in absence of food intake) and consumptive self-administration of food, and to determine if these effects are independent of aversive and metabolic effects. Using both operant and non-operant procedures, we examined the effect of microinjection of the mixed melanocortin receptor 3/4 agonist melanotan-II (MT-II) into the NAcc on both appetitive and consumptive behaviors.

## METHODS

### Animals

Adult, male, C57BL6/J mice (The Jackson Laboratory) were used for all studies and arrived at 2-3 months of age. Mice were housed in standard clear acrylic shoebox containers with ad libitum food (Purina Lab Chow, 5001) and water for at least 7 days to acclimate to the vivarium. The housing room was maintained on a 12:12-h light:dark cycle. All procedures were approved by the IACUC before the experiments were conducted. The principles of laboratory animal care and the National Research Council “Guidelines for the Care and Use of Mammals in Neuroscience and Behavioral Research” were followed by all laboratory personnel.

### Surgery and Microinjections

Mice were anesthetized with avertin (2, 2, 2-tribromoethanol, Sigma Aldrich, St. Louis, MO) or isoflurane for stereotaxic surgery (Model 1900 Stereotaxic Alignment System, 1-micron resolution, David Kopf Instruments, Tujunga, CA). To avoid piercing a ventricle, bilateral cannulae (26 g, stainless steel tubing, 10.0 mm in length) were placed on a 20-degree angle from perpendicular with the tip of the cannulae ending 1 mm above the NAcc (cannula entering at A/P =+1.650, M/L = +/-2.436, D/V =-3.6 relative to bregma). The cannulae were affixed to the skull using anchor screws (#000-120, Antrin Miniature Specialties) and Durelon cement (3M EPSE, St. Paul, MN). The mice were checked and handled regularly after surgery to monitor body weight, check integrity of the cannulae and stylets, and habituate the mice to mild restraint. Mice recovered for 1 week prior to their first microinjection. At the end of each study, the mice were euthanized, and the brains were collected to verify cannulae placements (see Figure 1 for cannulae locations for mice in the operant and conditioned taste avoidance studies).

**Figure 1:**
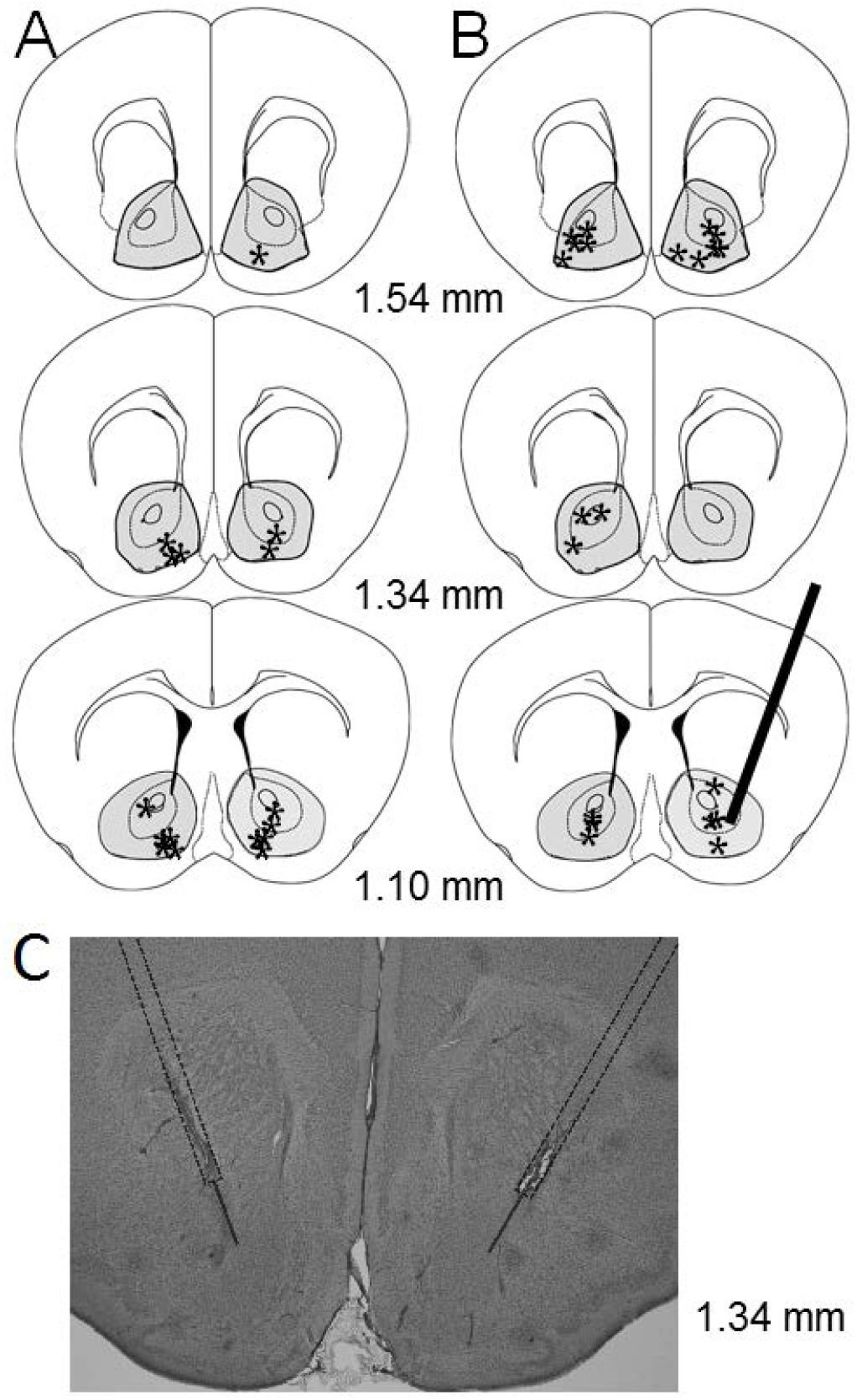
Cannulae placements in the nucleus accumbens. The locations of the cannulae placements for the operant study (panel A) and the CTA study (panel B) are indicated by asterisks. The distance from Bregma of the coronal slices is indicated between the panels. A hypothetical cannula tract (coming in at a 20-degree angle) is show in the bottom right slice of panel B. A coronal section of mouse brain showing a simulation of cannulae placement and injectors to demonstrate an acceptable area for injection into the NAcc (panel C).

Injections were conducted by restraining the mouse gently in one hand, removing the stylets (33 g stainless steel wire), and inserting the injector (33 g, stainless steel tubing, 11.0 mm in length) into the guide cannula. The injection of 100 nl was given over 60 seconds and the injector was left in place for an additional 30 seconds to allow diffusion of the bolus away from the injector. The stylet was then replaced, and the procedure was repeated on the other side.

### Home cage procedure

Approximately 16 hours before injection (2 hours before lights off), mice (n=12) were moved to a clean cage with no access to food. This short period of food deprivation enabled us to observe a decrease in feeding following injection of melanotan-II (MT-II) compared to control injection of saline. For injections, the mice were weighed and then injected bilaterally with either saline or MT-II before being placed back in their home cage with a pre-measured amount of food. The food was weighed again at 1, 2, 4, 6, and 24 hours after injection to determine how much food was consumed. All mice received injections of saline and each dose of MT-II (0.1, 0.3, and 1 nmol/side) for a total of 4 injections per mouse. The order of injection doses was counterbalanced between mice in the group, with at least 3 days between injections.

### Operant procedure

At least 7 days after surgery, mice (n=7) were moved into the operant chambers (Med Associates, Georgia, VT). Each operant chamber was contained in a sound attenuating exterior cabinet with a fan to mask external noise. A retractable lever was located on one wall of the chamber with a food magazine located immediately adjacent to the lever. The food hopper delivered 20 mg pellets of normal mouse chow diet (Research Diets, New Brunswick, NJ, USA). A metal sipper tube connected to an inverted conical tube containing water was located opposite the food hopper and was available throughout the experiment. Licks on the sipper tube were monitored and recorded through a lickometer (Med Associates, Georgia, VT). A house light was located above the food hopper and the mice were maintained on a 12-hour light/dark cycle with lights on at 7 am and off at 7 pm. Each mouse was provided with an acrylic square (approximately 5 cm^2^) in the chamber to provide an alternative surface to the metal rod floor for a sleeping perch. Operant training was similar to that previously published (Richard et al., 2011). Mice were required to complete 10 lever presses for delivery of each 20-mg pellet on the first day (fixed ratio 10, FR10). Following successful completion of the FR10, the lever retracted from the chamber for 10 seconds and the pellet was delivered into the food magazine. After 10 seconds, the lever was re-extended into the chamber and the mouse was allowed to respond for additional pellets. On the second and subsequent days, the FR was increased to FR30 for each pellet with a 10 second timeout after completion of the FR. Mice had access to the lever and water at all times during the study when they were in the chambers. Mice were only removed from the chambers for routine cleaning, restarting of the session, and for injections and handling. After at least a week to acclimate to the operant chambers, the effect of MT-II on operant responding for food was assessed.

Approximately 1 hr before lights out on injection days, the mice were removed from their chambers, weighed, and then 100 nl of the vehicle (sterile saline) or MT-II (0.3 nmol) was injected bilaterally as previously described. Following injection on both sides, the stylets were replaced, and the mouse was placed back into the holding cage for either 15 minutes (consumptive sessions) or 60 minutes (appetitive/non-reinforced responding sessions) before being returned to the operant chamber and the session begun. The effect of MT-II on responding for food was conducted first, with each mouse getting both saline and MT-II (order was counterbalanced between mice). For consumptive sessions the mice were on a FR30 as usual for 23 hr after the injection of MT-II. Lever pressing, the pattern of pellet delivery, and the pattern of water consumption over this time was monitored by Med-PC software. The data were then analyzed to determine the amount of food consumed 1, 2, 4, 6, and 23 hours into the session after the bilateral microinjection.

To determine the appetitive motivation to gain access to food in absence of eating, each mouse was placed back in the operant chamber 60 minutes after injection of saline or MT-II (again, order was counterbalanced) and the session was begun as normal except that the responding on the lever was not reinforced (no pellet was delivered and the lever did not retract following 30 responses) for 60 minutes. During this time the number and pattern of lever presses were recorded for analysis. After 60 minutes, the doors to the operant chamber enclosure were opened briefly, and the session was restarted on the usual FR30 for each pellet. Microinjections for the consumptive sessions were conducted approximately 3 days apart, and for the appetitive/non-reinforced sessions they were conducted with 7 days between injections. Each mouse received both vehicle and MT-II for both the consumptive and appetitive sessions, resulting in 4 injections per mouse.

### Conditioned Taste Avoidance (CTA)

After surgery to implant cannulae, mice were individually housed for at least 7 days prior to the beginning of the experiment with food and water available ad libitum. CTA procedures were similar to previously published methods (Lessov et al. 2001; Risinger and Cunningham 1998; Sharpe et al. 2005). Mice were then acclimated to limited (2 h) water access (approximately 10 am to noon) for at least 5 days before the start of the injections. Mice were weighed each day immediately preceding the limited access to fluid. Subjects were divided into two groups (saline vs. 1 nmol MT-II) based on daily water intake during the limited access period (n=6, saline; n=5, MT-II). A sodium chloride solution (0.2 M NaCl) was used as the novel flavor that was paired with the microinjection of MT-II or vehicle. The NaCl solution was presented every other day, immediately after the mice were weighed and was only available for 1 hour. With the exception of the first exposure (D1) and the last exposure (D11), mice were immediately microinjected with MT-II after the bottle containing the NaCl was removed from the cage. Data from the first day of NaCl exposure was not used to minimize bias due to any neophobia to the solution on the first day. For data analysis, consumption of NaCl before the first pairing was used as baseline NaCl consumption and was compared to NaCl intake on the 4 days following MT-II pairing with NaCl. To prevent dehydration, on days that NaCl was given water was made available to each mouse for 30 minutes beginning 4 hours after NaCl was removed. Water was available as usual (for 2 h) on intervening days that NaCl was not presented.

### Indirect calorimetry

Mice (n = 7, male C57BL6/J) with bilateral cannuale placed above the NAcc, as described above, were habituated to the indirect calorimetry chambers (OxyMax, Columbus Instruments). Mice received MT-II (0, 0.1, or 0.3 nmol/100 nl, bilaterally) in a counterbalanced order with at least 3 days between injections. A 2-hour baseline measure was taken in the OxyMax chambers before mice were briefly removed for microinjection and then replaced into the chamber for 5 hours of recording. The sample air from each chamber was analyzed for O_2_ and CO_2_ to determine O_2_ and CO_2_ content. From these measures, oxygen consumption (vO_2_), heat production, and respiratory exchange ratio (RER) were measured over this 7-hour span (2 hours before injection and 5 hours afterwards). The 5 hours after MT-II injection was chosen to include the time period for the anorexic effects of MT-II in the other studies. These measures were taken in the dark portion of the light cycle (similar to the feeding studies) with food and water available ad libitum during the time in the OxyMax chambers. Mice were euthanized at the end of the study and cannulae placement was confirmed histologically, with 100% of the mice having accurate cannulae placement.

### Drugs

MT-II (Phoenix Pharmaceuticals, Burlingame, CA) was reconstituted in sterile saline in a stock solution at a concentration of 10 nmol/ul and stored at -20 C. The stock solution was diluted with sterile saline as needed to prepare the lower concentrations of MT-II for injection.

### Statistics

Both the home cage and operant intake data were analyzed initially by repeated-measures ANOVA (RM-ANOVA) with cumulative food intake as the independent variable, and time post injection and MT-II dose as the dependent variables. To determine if there were significant differences between the doses tested from vehicle, significant interactions between time and dose were followed up with either RM-ANOVA at each dose (with post-hoc Dunnett test) for the home cage data, or with planned comparison between the two doses at each time point for the operant data. The CTA data were analyzed by RM-AVOVA with NaCl consumption during the 1- h limited access (in ml) as the independent variable and with pairings as the within-subject repeated measure and drug (vehicle or MT-II) as the between subjects dependent variables. *P* < 0.05 was considered significant.

## RESULTS

### Home cage Study

MT-II injection into the NAcc decreased food consumption, with a significant main effect of drug (*P* < 0.0001, F_3,33_ = 12.7), time (*P* < 0.0001, F_4,44_ = 1008.5), and a significant interaction between drug and time (*P* < 0.004, F_12,132_ = 2.6). All doses of MT-II significantly decreased food intake at 1, 2, and 4 hours after injection compared to saline (Figure 2). By 24-h after injection of MT-II, only the 0.3 nmol MT-II dose was still significantly decreased compared to intake after vehicle injection.

**Figure 2:**
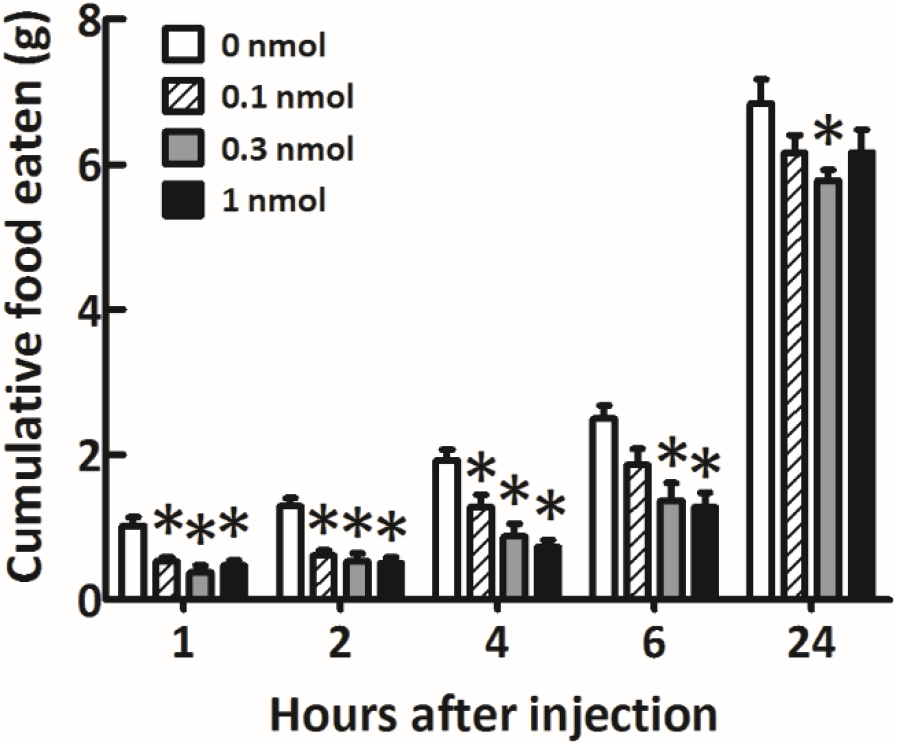
MT-II decreases food intake in the home cage. **S**ixteen-hour food deprived mice were injected bilaterally into the NAcc with MT-II in a repeated measures design. Every dose of MT-II tested produced a significant decrease in food intake at 1, 2, 4, and 6 hours, with only the effect at 0.3 nmol persisting until 24 hours post injection. * *P* < 0.05 compared to saline at same time point.

### Operant Study

Similar to results in the free access home cage study, intra-accumbens injection of MT-II decreased responding for food in an operant paradigm. There was a significant main effect of time (*P* < 0.0001, F_4,24_ = 123.8) but not treatment (*P* = 0.28). However, there was a significant interaction between time and treatment (*P* = 0.0005, F_4,24_ = 7.4). Planned comparison analysis at each time point corresponding to the home cage study revealed a significant decrease in pellet delivery at 2 and 4 hours post injection of MT-II compared to vehicle (Figure 3A). Representative cumulative records from one mouse following MT-II and saline injections show the disrupted lever pressing for food pellets in the first 4 hours following injection of MT-II compared to saline treatment day (Figure 3B and C).

**Fig 3:**
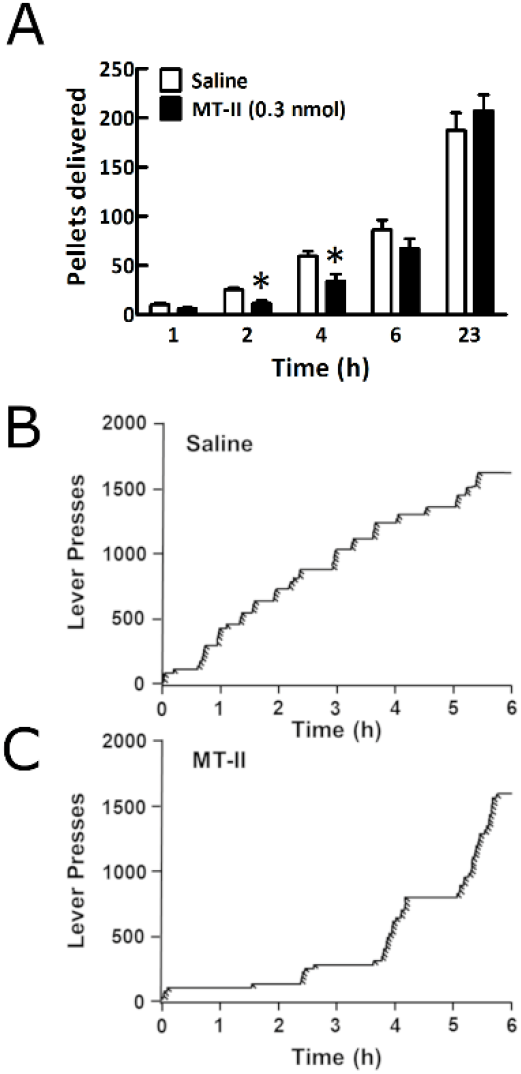
Operant responding for food is decreased following MT-II. Mice were injected with saline or MT-II (0.3 nmol) bilaterally into the NAcc one hour before the onset of the dark cycle (and 15 minutes before the start of the operant session). In a within animal design, MT-II decreased responding for food pellets compared to saline at 2 and 4 hours into the operant session (A). Sample cumulative records for operant behavior from the same mouse following saline (B) and MT-II (C) injection show that responding for food is disrupted at the beginning of the session. Near normal responding for food resumes approximately 4 hours into the operant session following MT- II. Vertical deflection of the line depicts lever presses and downward slashes indicate delivery of a food pellet. * *P* < 0.05 compared to saline at same time point.

In the appetitive study with 60-minutes of non-reinforced responding, lever pressing (appetitive responding) was decreased by MT-II with a significant main effect of drug (*P* = 0.01, F1,6 = 12.1) and time (*P* < 0.0001, F8,48 = 36.5), and interaction between drug and time (*P* = 0.006, F8,48 = 3.2). Planned comparisons between MT-II and vehicle at various time intervals (1, 2, 5, 10, 20, 30, 40, 50 and 60 minutes) showed significant (P < 0.05) decreases in cumulative lever presses at all time points except for 30 and 40 minutes (*P* < 0.06) (Figure 4A). Cumulative records of non-reinforced lever pressing from one representative mouse following injection of both saline and MT-II demonstrate the interruption in appetitive responding caused by injection of the melanocortin receptor agonist in the NAcc (Figure 4B and C).

**Figure 4:**
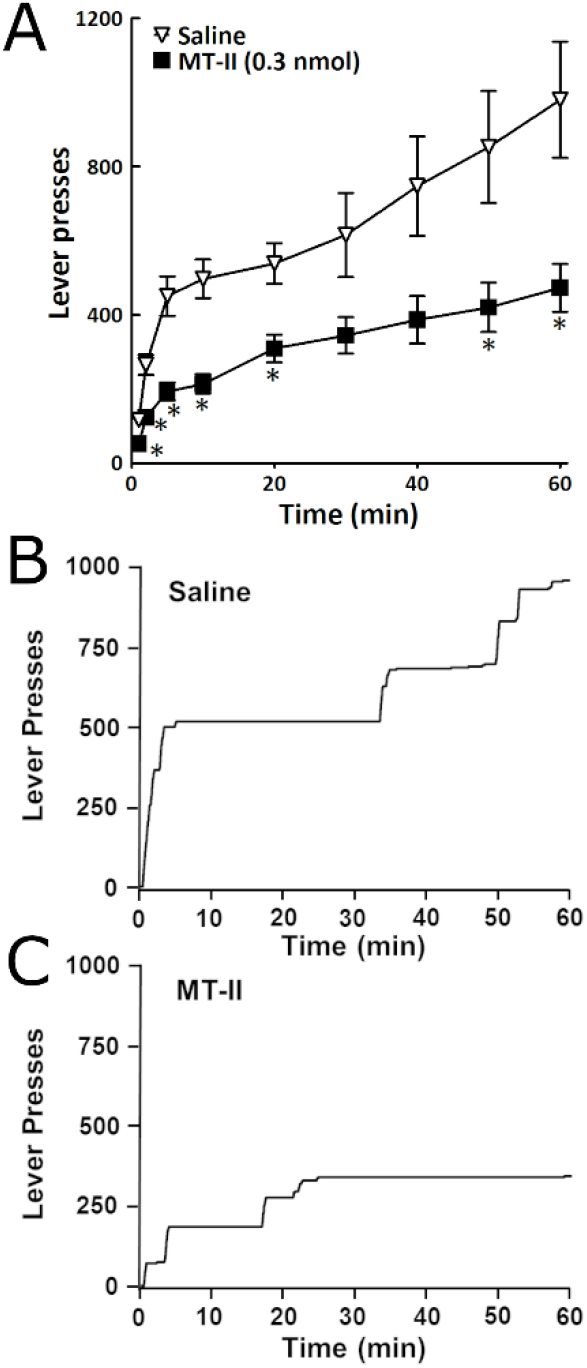
Appetitive responding for food is decreased by MT-II in the nucleus accumbens. Mice were placed in the operant chamber 60 minutes following injection of MT-II (0.3 nmol) into the NAcc and allowed to respond on the lever for 60 minutes during which time no food pellets were delivered. Non-reinforced (or extinction) responding was used as a measure of appetitive motivation. MT-II significantly decreased non-reinforced lever pressing compared to saline injection (A). A sample cumulative record from one mouse following saline (B) and MT-II (C) injection shows that although the initial rate of responding was similar between the treatments, the responding after MT-II was not maintained and remained depressed throughout the 60-minute test. Vertical deflection in the line is indicative of lever pressing. * *P* < 0.05 compared to saline at that time point.

### Conditioned Taste Avoidance

Using a procedure that has produced a significant conditioned taste avoidance previously in response to a high dose of ethanol (Sharpe et al. 2005), there was no difference in NaCl consumption between mice injected with MT-II versus vehicle (main effect of treatment, *P* = 0.83) (Figure 5). While food intake was not measured following MT-II injection in this experiment, there was a significant drop in body weight 24-h after injection when compared to vehicle injected mice (data not shown), consistent with the decreased food intake caused by the same dose of MT-II in the home cage study (see Figure 2).

**Figure 5:**
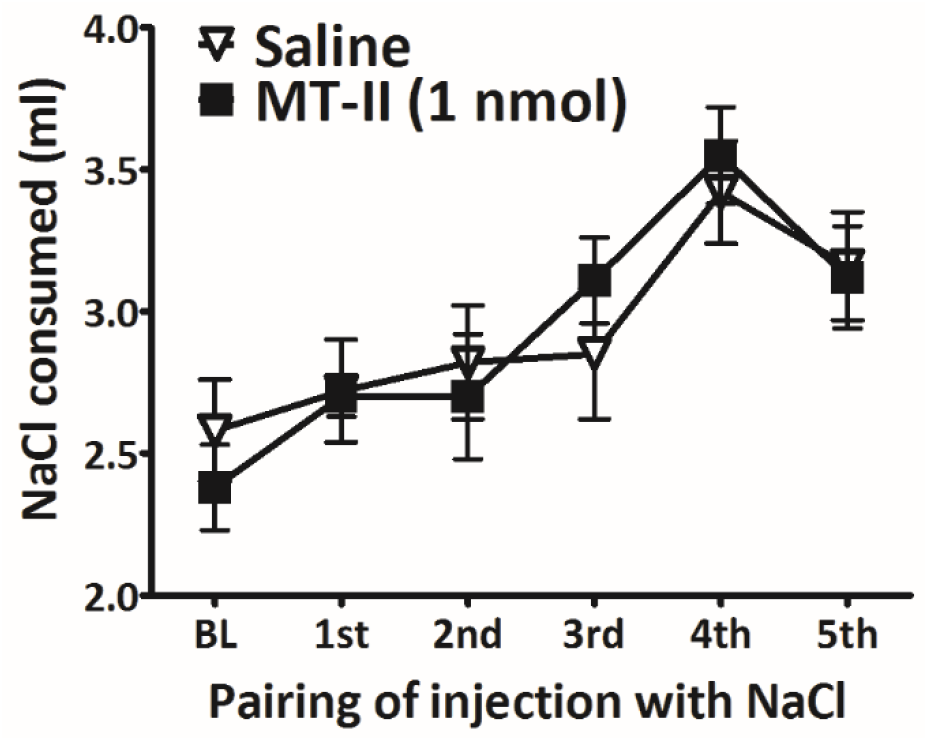
MT-II injected into the nucleus accumbens does not condition a taste avoidance. The highest dose of MT-II tested in the original home cage feeding study (1 nmol) did not condition a taste avoidance following 5 pairings of MT-II with a novel NaCl solution. Fluid restricted (2-h access to water/day) mice did not decrease intake of the NaCl solution that was paired with intra-accumbens injection of MT-II, suggesting that MT-II did not produce aversive stimuli.

### Indirect Calorimetry

Data from the indirect calorimetry were averaged over 30-minute bins for the entire 7-hour testing period (2 hours before microinjection, 5 hours afterwards). Following MT-II injection into the NAcc there were no significant changes in RER, vO2, or heat production from saline at any time point measured (Figure 6).

**Figure 6:**
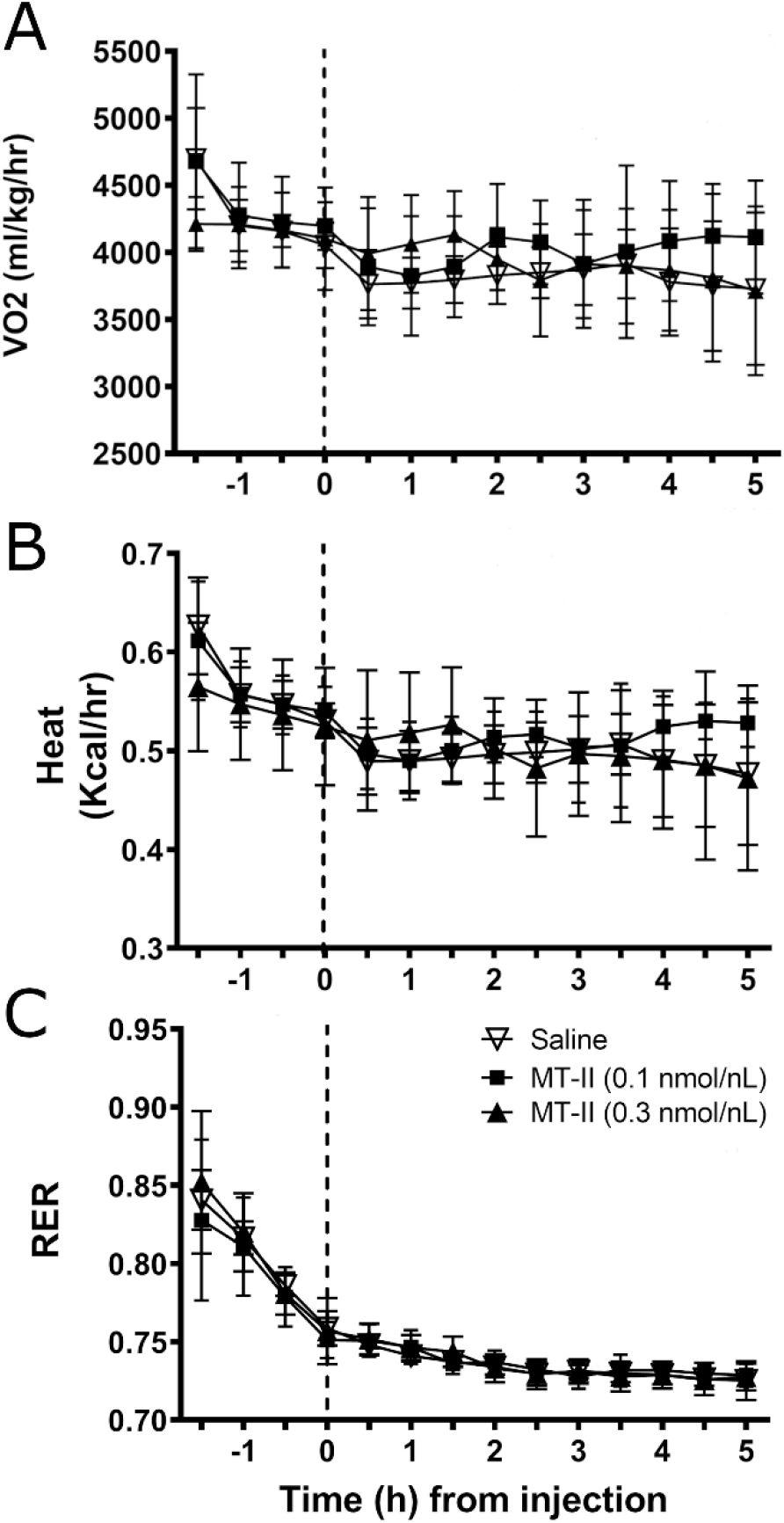
MT-II injected into the nucleus accumbens does not affect measures of metabolism measured by indirect calorimetry. In the same timescale that MT-II decreases feeding and appetitive responding it has no effect on oxygen consumption (VO2) (panel A), heat production (panel B), or calculated respiratory exchange ratio (RER).

## DISCUSSION

Numerous previous studies have investigated the role of melanocortins in the amygdala, midbrain, brainstem, and various regions of the hypothalamus on feeding and reward (Dunigan & Roseberry, 2022; Skibicka and Grill 2009; Wirth et al. 2001). However, POMC neurons in the arcuate nucleus of the hypothalamus also project to the NAcc (Bagnol et al. 1999; Kitahama et al. 1986; Lim et al., 2012), an area of the brain that is highly implicated in reward. These projections, and their associated melanocortin receptors, are present in the NAcc (Mountjoy et al. 1994; Pandit et al., 2015). There is a physiological role of melanocortins in feeding and sucrose intake in the NAcc (Pandit et al., 2015; Lerma-Cabrera et al., 2012), but prior studies did not examine if melanocortin agonists decrease intake via aversive properties or have any effect on metabolism.

Our data extend previous findings in rats by demonstrating in ad libitum fed mice that administration of a melanocortin receptor agonist in the NAcc decreases both appetitive and consumptive food behaviors for rodent chow without eliciting aversive or metabolic properties. Injection of the melanocortin receptor agonist MT-II bilaterally into the NAcc decreased food consumption in both home cage (free feeding) and operant self-administration, with the greatest effect seen 1-4 hours after injection. In addition, MT-II decreased non-reinforced responding for food, suggesting that appetitive drive or motivation to eat is also decreased by melanocortin receptor agonists in the NAcc. However, these effects are not mediated by general aversive properties since microinjection with the highest dose of MT-II tested was unable to produce aversive behaviors in a conditioned taste study. Additionally, although MT-II administered into the hypothalamus or ventricles increases metabolism, injection into the NAcc had no effect on metabolism measured by indirect calorimetry. This suggests that melanocortins can decrease appetite or motivation to access food independent of post-ingestive properties of the food, such as palatability. Factors that influence the initiation and maintenance of food consumption are food availability, food palatability, experience, metabolic need (hunger), and satiety, among others.

One possible interpretation of our results is that melanocortin receptor agonists decrease both the appetitive and consumptive behaviors through a general decrease in the perceived value of the food. This could be elicited by a decrease in perceived need for food—a decrease in hunger, an increase in satiety, or ambivalence to eating. While Pandit et al. (2015) examined the effect of intra-accumbens injection of melanocortin agonist on progressive ratio in rats to obtain sucrose, progressive ratio testing procedurally mixes reward consumption with measure of motivation to obtain the reinforcer. This measure is, by default, not a pure measure of motivation in absence of intake since a shift in satiety would be confounded by the consumption during responding.

Similar effects on appetitive and consumptive behaviors to what we report here have been reported following pre-feeding (Azarra et al. 2002; Ishii et al. 2003) as well as after injection of cannabinoid antagonists (Thornton-Jones et al. 2005). These studies suggested that pre-feeding resulted in a decrease in motivation to eat and a premature transition to satiety sequence as opposed to conditions of no pre-feeding or cannabinoid antagonist. While we did not examine the satiety sequence following MT-II versus vehicle treatment in our model, a shift towards satiety following MT-II injection would be consistent with the results that we observed. This hypothesis agrees with a previous study examining the effect of <u>ventricular</u> MT-II on feeding microstructure which concluded that MT-II decreased feeding primarily by affecting satiety, and not by inducing taste deficits (Williams et al. 2002).

Although an impairment in locomotor function could also decrease responding for and consumption of food, the rate of responses made on the lever for food pellets in our studies was similar on MT-II treated days compared to vehicle treatment days (for an example, see Figs. 3 panels B & C and 4 panels B & C). Although we did not specifically examine locomotor activity in these subjects in response to MT-II, other groups have reported no decrease in locomotor activity in mice following ventricular injection of MT-II in rats (Fan et al., 1997; Williams et al., 2002).

In these studies, we took several measures to increase confidence in the site-specificity of the results. The number of microinjections per mouse was limited to decrease damage that could be produced by multiple injections. In addition, we used a 20-degree angle of the cannulae, small injection volume (100 nl), slow injection rate, and delayed removal of the injector specifically to limit the possibility of diffusion outside the NAcc or entering the ventricle. The lack of metabolic effects of MT-II certainly support that there was no leak of MT-II into the ventricle.

Future studies should be conducted to test the hypothesis that melanocortin receptor agonists in the NAcc devalue the reinforcer and shift the animal into a sated state. Furthermore, it should be addressed that these studies, and those preceding it, were all completed with only melanocortin peptides to mimic therapeutic treatment. However, endogenous POMC stimulation likely results in much lower peptide levels in addition to co-release of POMC-derived peptides. Beta-endorphin is another POMC peptide product that is an agonist at mu-opioid receptors and is believed to be co-released with alpha-MSH. However, in contrast to melanocortin receptor agonists, mu-opioid receptor agonist delivery in the NAcc <u>increases</u> food intake (Mucha & Iversen, 1986; Peciña & Berridge, 2000). As these POMC-derived neuropeptides are co-released and have opposite effects on feeding, the appetitive field would greatly benefit from a deeper understanding of how stimulation of POMC projections to NAcc modulate reward learning and consumption. We suggest from our results that melanocortin projections to the NAcc regulate motivation to eat as well as the amount of food that is consumed when food is available. Development of a therapeutic capable of shifting an organism towards a state of satiety, thus decreasing food consumption and motivation to eat would be useful in the treatment of obesity.

## Acknowledgements

The authors would like to thank Veronica Otero-Corchon for her assistance with the conditioned taste avoidance study. This work was supported by the following grants: National Institutes of Health grant DK066604 to MJL, P20GM125528 (sub-project 5338 to ALS), and SC3GM121214 to ALS; American Heart Association National Scientist Development grant to ALS; and Presbyterian Health Foundation Seed Grants to ALS. The authors were solely responsible for the content.

